# *rTPC* and *nls.multstart*: a new pipeline to fit thermal performance curves in *R*

**DOI:** 10.1101/2020.12.16.423089

**Authors:** Daniel Padfield, Hannah O’Sullivan, Samraat Pawar

## Abstract

1. The quantification of thermal performance curves (TPCs) for biological rates has many applications to problems such as predicting species’ responses to climate change. There is currently no widely used open-source pipeline to fit mathematical TPC models to data, which limits the transparency and reproducibility of the curve fitting process underlying applications of TPCs.
2. We present a new pipeline in *R* that currently allows for reproducible fitting of 24 different TPC models using non-linear least squares (NLLS) regression. The pipeline consists of two packages – *rTPC* and *nls. multstart* – that allow multiple start values for NLLS fitting and provides helper functions for setting start parameters. This pipeline overcomes previous problems that have made NLLS fitting and estimation of key parameters difficult or unreliable.
3. We demonstrate how *rTPC* and *nls*.*multstart* can be combined with other packages in *R* to robustly and reproducibly fit multiple models to multiple TPC datasets at once. In addition, we show how model selection or averaging, weighted model fitting, and bootstrapping can easily be implemented within the pipeline.
4. This new pipeline provides a flexible and reproducible approach that makes the challenging task of fitting multiple TPC models to data accessible to a wide range of users.

## 1. Introduction

Thermal performance curves (TPCs) describe how biological rates such as growth, photosynthesis and respiration change with temperature. TPCs (and the parameters that underpin them) have been used widely in biology, from studying thermal adaptation (Schaum et al., 2017; Smith et al., 2019), to predicting ectotherm range shifts (Sunday, Bates, & Dulvy, 2012; Sinclair et al., 2016) and changes in disease dynamics (Molnár, Kutz, Hoar, & Dobson, 2013; Cohen et al., 2017; Mordecai et al., 2019) under expected climate change. Despite their wide use across ecology and evolution, there is no open-source, flexible approach available to fit TPC models to data (henceforth simply “fit TPCs”) that allows for reproducible fitting using NLLS. Current software for fitting TPCs, such as the *R* packages *temperatureresponse* (Low-Décarie et al., 2017) and *devRate* (Rebaudo, Struelens, & Dangles, 2018), do not address the well-known sensitivity of NLLS algorithms to parameter starting values, which is exacerbated when fitting multiple models with varying non-linearities, and to multiple datasets with differences in sampling, rate measurements and coverage of temperature ranges. Moreover, these existing packages do not address robust quantification of parameter uncertainty.

Many different mathematical models have been used to fit TPCs (Krenek, Berendonk, & Petzoldt, 2011; DeLong et al., 2017; Low-Décarie et al., 2017) which can make it difficult to determine the “best” model for any given dataset. A few papers have evaluated the performance of TPC models (Angilletta Jr, 2006; Shi & Ge, 2010; Krenek et al., 2011). The most comprehensive analysis to date compared 12 models, and demonstrated how model choice alters the predicted species-level response to temperature (Low-Décarie et al., 2017). However, using model selection to select the best or most appropriate model for specific datasets remains rare (but see Montagnes, Morgan, Bissinger, Atkinson, & Weisse, 2008). Instead, a single model is used, often chosen for its mechanistic underpinnings, despite little agreement about the mechanistic links between enzyme kinetics and emergent biological rates (e.g. velocity, feeding rate, growth rate). Others prefer to use a model that directly estimates desired parameters (e.g. optimum temperature). There is likely no “best” model to use for fitting TPCs, with different models proving the most appropriate for different biological processes, taxa, and levels of data quality. Consequently, a new analysis pipeline that allows users to fit TPCs, while remaining flexible to the research question being asked, is sorely needed.

Here, we present *rTPC* and *nls*.*multstart*; two open-source *R* packages that provide the basis for a pipeline to robustly and reproducibly fit TPCs. The pipeline allows the fitting of 24 different TPC model formulations, and we demonstrate how multiple models can be fitted to the same curve, as well as how multiple datasets can be fitted. We also describe helper functions within *rTPC* for the estimation of start parameters, upper and lower parameter limits, and commonly-used parameters (e.g. optimum temperature, activation energy or Q10). Finally, we illustrate how this pipeline can be used for model selection and model averaging, as well as how weighted model fitting and bootstrapping implemented using *rTPC* can be used to account for parameter and model uncertainty.

## 2. Pipeline overview

The goal of *rTPC* and the associated pipeline is to make fitting TPCs easier and repeatable. Extensive examples of the pipeline can be found at https://padpadpadpad.github.io/rTPC where all vignettes are available. When developing *rTPC*, we made a conscious decision not to repeat code and methods that are already optimised and available in the *R* ecosystem. Instead, they are utilised and incorporated into the pipeline.

### 2.1 Models contained in *rTPC*

*rTPC* contains 24 mathematical TPC models (Table S1). Most models are named after the author that first formulated the model and the year of its first use (e.g. *thomas_2012()*). A list of all models in *rTPC* can be accessed using *get_model_names()*. Models can be characterised by whether they appropriately model negative rates before and after the optimum temperature (Table S1).

### 2.2 NLLS fitting using multiple start parameters using *nls*.*multstart*

The Gauss-Newton (implemented in *nls*) and the Levenberg-Marquardt (implemented in *minpack*.*lm::nlsLM*) NLLS fitting algorithms are sensitive to the choice of starting values for the model parameters. This sensitivity can result in differences in parameter estimates between separate fitting attempts for the same dataset, or a complete failure to fit the model (the optimisation does not converge). To address this, the *R* package *nls*.*multstart* – and its only function *nls_multstart()* - generates multiple start values and fits many iterations of the model using the Levenberg-Marquardt algorithm implemented in *nlsLM*. The best model is then picked and returned using Akaike’s Information Criterion corrected for small sample size (AICc) (Padfield & Matheson, 2018).

### 2.3 Estimating starting parameter values and limits for fitting TPCs using *rTPC*

*rTPC* helper functions *get_start_vals(), get_lower()* and *get_upper()* aid in the specification of sensible start values and limits that can be used by *nls_multstart()* (or *nls* and *nlsLM*). These functions return values for the desired model, which is specified using the argument *model_name*. Where possible, the model’s starting parameter values are estimated from the data. In all other instances, start values are the average fitted parameters from studies that used that equation. Upper and lower limits are set at biologically implausible values. If these helper functions aren’t needed, values can be set manually.

### 2.4 Calculating derived TPC parameters

Parameters of TPCs (such as optimum temperature or Q10) are commonly used in downstream analyses (e.g. to determine if optimum temperature correlates with local climate across taxa). However, the best-fitting model’s parameters may not include the parameter of interest. *rTPC* has helper functions, such as *get_topt()* and *get_rmax()*, that numerically calculate most parameters of interest from any fitted TPC (Table 1). These derived parameters are calculated from high resolution predictions (0.001°C intervals) of the fitted model. The function *calc_params()* returns values for all 11 derived parameters in a dataframe (Table 1). *calc_params()* does not return estimates of uncertainty in these derived parameters, which we address below (section 3.3).

**Table 1:**
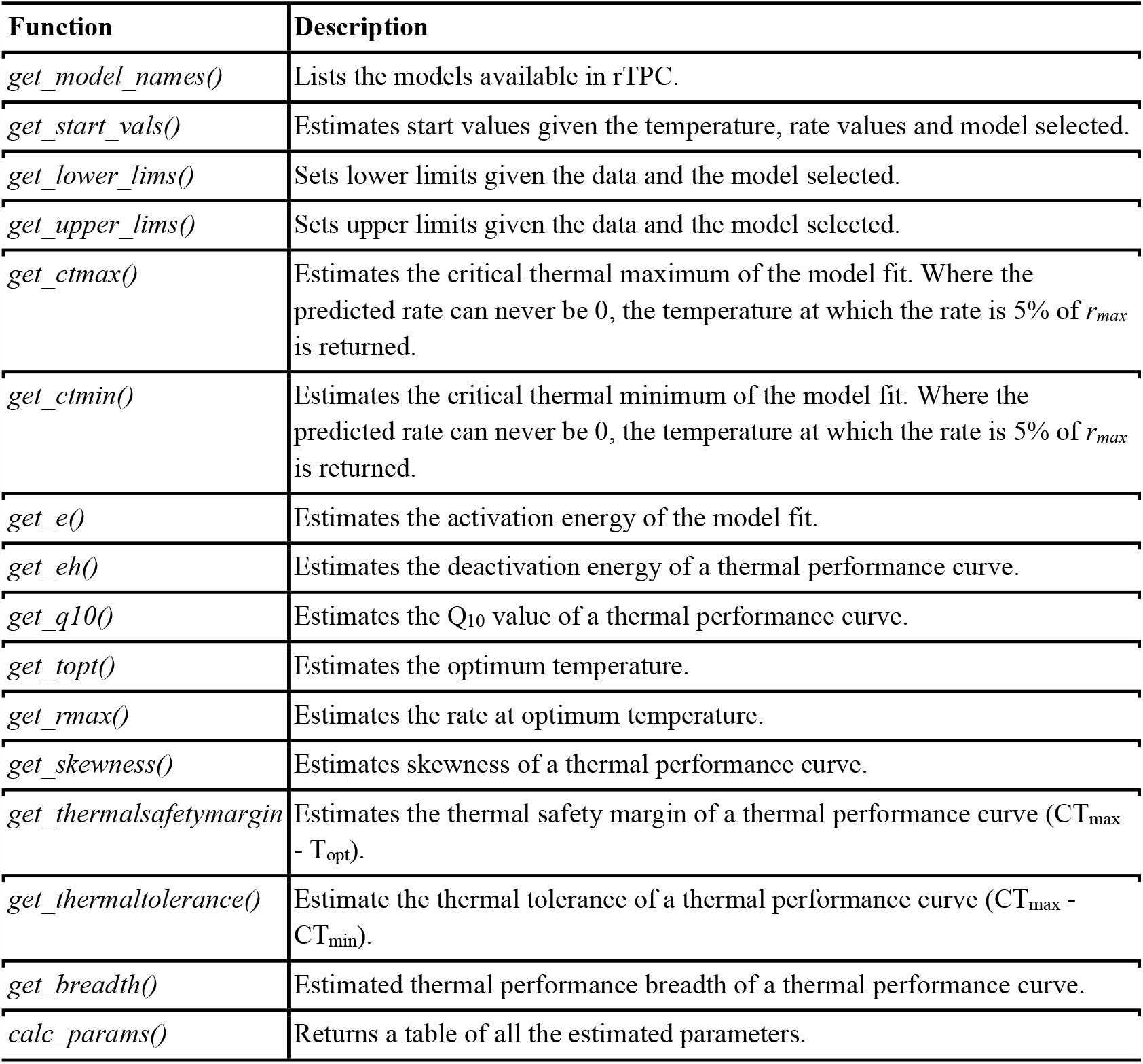
Overview of helper functions included in *rTPC*.

## 3. Uses for the *rTPC* pipeline

Below we give examples of potential applications and extensions to the pipeline, why they are important, and guidance as to how they can be incorporated.

### 3.1 Model selection and model averaging

The “best” model for one dataset is not necessarily the best across other datasets. Our pipeline provides a flexible approach to help with model selection. For example, after fitting a number of potential models, AICc scores can be used to rank the models for each individual curve fit and pick the best overall model across all curves in a dataset. Alternatively, one may choose the best model specific to each TPC, or use model averaging to obtain an overall TPC curve and parameter estimates by weighting each model’s fit by its AICc (Figure 2). *vignette(“model_selection_averaging”)* provides an example of how to implement model selection and model averaging.

**Figure 1.**
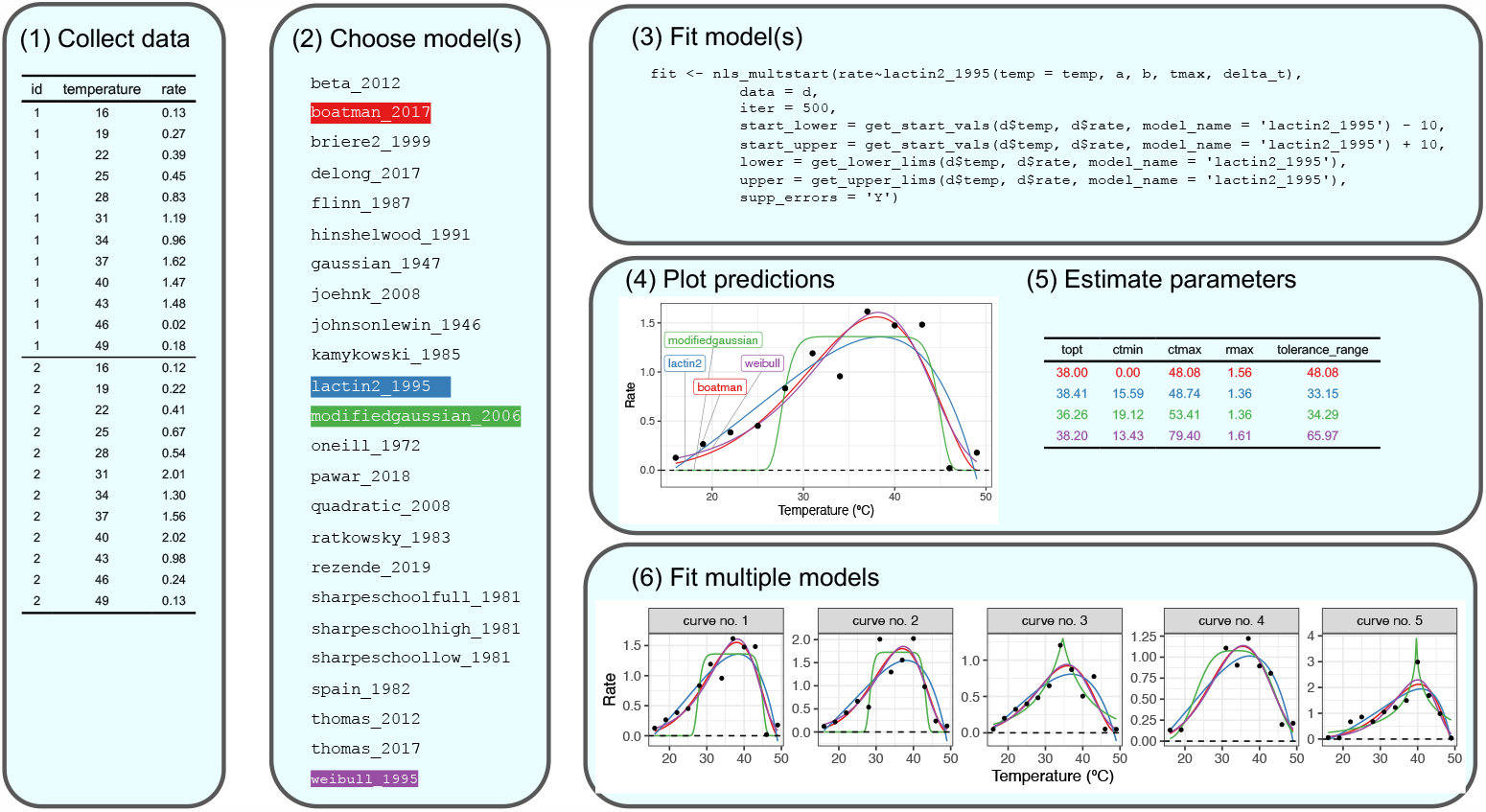
General pipeline for fitting thermal performance curves using *rTPC*. (1) Collect, check, and present data in long format. (2) Choose which models from *rTPC* to be use. Here, a random assortment of four models were chosen. (3) Fit the models using *nls*.*multstart* and helper functions from *rTPC*. (4) Models can be visualised and (5) common traits of TPCs can be estimated using *rTPC::calc_params()*. (6) This simple pipeline can be scaled up to be used on multiple curves.

**Figure 2.**
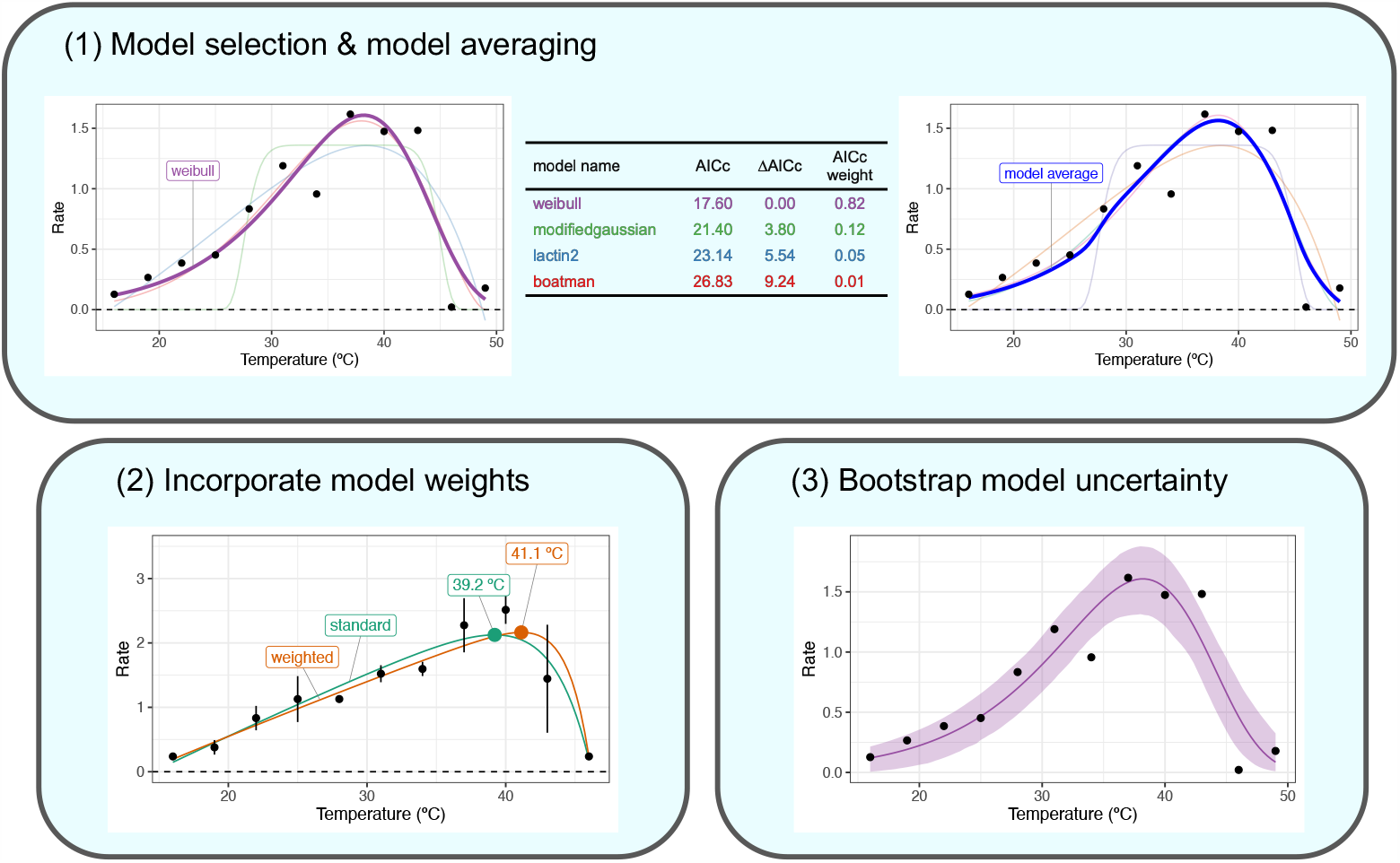
Potential applications for fitting thermal performance curves using *rTPC*. (1) AICc scores of model fits can be calculated to help with model selection or model averaging. (2) If TPCs are fit to averages of replicates, weighted NLLS can be used to reduce parameter bias. (3) After the model has been fitted, non-parametric bootstrapping can estimate model uncertainty and confidence intervals for parameters.

### 3.2 Data-weighted TPC model fitting

Due to non-independence of replicate rate measurements across temperatures, the mean rate at each temperature is often taken before fitting the TPC. This approach ignores variation in measurement values at each temperature. Incorporating variation in measurement errors across temperatures can <u>be</u> essential to improving the model fit and reducing biases in parameter estimates (Figure 2) (Davison & Hinkley, 1997). This can be implemented using weighted NLLS fitting, which can be applied using most methods of fitting NLLS in *R*. The optimal way to apply weights is to use *1/standard deviation*, which must be included as a vector the same length as the sample size. *vignette(“model_weighting”)* provides an example of how to implement weighted NLLS when fitting TPCs.

### 3.3 Quantifying uncertainty in model fits and parameter estimates

Quantifying uncertainty in the TPC model fit as a whole and the estimated parameters is challenging. The recommended method is to calculate confidence intervals around model parameters of the TPC by constructing the likelihood profile of the parameters (most commonly done by using *stats::confint* which invokes *MASS::confint*.*nls* or *nlstools::confint2* (Baty et al., 2015)). However, in many instances the profiling (a numerical method) does not converge or the likelihood profile that emerges is asymmetric or skewed. Consequently, parameter and model prediction confidence intervals of TPCs calculated in this way can be unreliable. Moreover, although profiling returns confidence intervals of the model parameters, this method cannot calculate the uncertainty in derived parameters (section 2.4).

Bootstrapping is a robust alternative to computing both the parameter and model prediction confidence intervals. Non-parametric bootstrapping entails resampling a dataset repeatedly and re-fitting the model to reconstruct a relatively unbiased sampling distribution of the parameters. Parameter confidence bounds can then be constructed using this distribution. Bootstrapping can also be used to calculate confidence intervals of derived parameters. The *rTPC* workflow uses the *Boot()* function from the *R* package *car* (Fox, 2006), which implements two types of non-parametric bootstrapping: case and residual resampling. In case resampling the data themselves are sampled (with replacement) to create a distribution of resampled parameter estimates, while residual bootstrapping uses mean centred residuals to create the distribution. Both methods have their pros and cons, and we leave it to the user to decide which one to use (or evaluate their performance for fitting TPCs). Non-overlapping confidence intervals of parameters between different TPCs may be used for inference, but these should be treated with caution for TPCs as datasets are often too small, making this type of inference unreliable. *vignette(“bootstrapping_models”)* provides an example of how to implement bootstrapping for TPC models using *rTPC* and *car::Boot()*.

Finally, data-weighted TPC fitting (section 3.2) can be combined with bootstrapping to potentially yield both unbiased parameter estimates and better estimates of uncertainty. *car::Boot()* now supports both case and residual resampling for weighted NLLS and *vignette(“weighted_bootstrapping”)* provides an example of how to implement this when fitting TPCs.

## 4. Key considerations when fitting TPCs

Effective fitting of TPCs depends on decisions made during experimental design, data collection, and model choice.

### 4.1 Data considerations

For effective fitting of TPCs, the number of unique temperature values used, the level of replication at each temperature, and the temperature range, all need to be considered. In the (common) scenario where all three cannot be maximised, the objective of the TPC fitting - and the parameters of particular interest - need to be considered. For example, in thermodynamic models, if the objective is to quantify the activation energy accurately, thermal range can be traded off for a finer degree of temperature resolution in the operational temperature range of the study organism (Pawar, Dell, Savage, & Knies, 2016). It is particularly important to consider the level of replication at each temperature: sampling multiple individuals at each temperature can give multiple individual TPCs of a population.

### 4.2 Which models to fit

The decision on which TPC models to fit largely depends on the type and quality of data, and the questions being asked. In terms of the data, there need to be at least *k* + 1 points for fitting a model, where *k* is the number of model parameters. However, in NLLS fitting, the minimum number of data points needed to reliably fit a model to data can vary with the mathematical structure of the model (Burnham & Anderson, 2002), so in general, “the more the merrier”. Carefully consider what model(s) you want to use before starting the analysis. If there are negative rate values, it is wise to fit models that can cross the x-axis both below and above the optimum temperature, such as *thomas_2012(), thomas_2017()* or *joehnk_2008()* (Table S1). In terms of the questions being asked, if there are specific traits of interest (e.g. optimum temperature), it may be beneficial to only consider models that explicitly include that parameter in their formulation. This may be especially pertinent for the activation energy, deactivation energy, and Q_10_, as they are sensitive to the calculation of the optimum temperature when calculated from model predictions. Finally, because NLLS is a numerical (inexact) model fitting method, consider carefully the correlations and mathematical relationships between parameters that may result in spurious parameter estimates (e.g. Kontopoulos, García-Carreras, Sal, Smith, & Pawar, 2018 in the case of the Sharpe-Schoolfield model)

## 5. Concluding remarks and future improvements

The pipeline presented here allows the user to fit TPCs in a simple, reproducible, and flexible framework. *rTPC* includes 24 model formulations previously used in the literature and *nls*.*multstart* provides a reliable method to fit non-linear models using multiple start values. It is important to note, however, that while this pipeline improves the fitting of TPCs, model fitting cannot fix poor data. In many experimental studies, the ideal approach to analysing TPCs would be with non-linear mixed effect models. This can be done using the *R* package *nlme* (Oddi, Miguez, Ghermandi, Bianchi, & Garibaldi, 2019), but Bayesian approaches have quickly become the easiest way to fit these types of models in *R*. Additional functionality of *rTPC* would be to output formatted code of the model equation, start parameters, and parameter limits that could be used by *brms*, a package that fits Bayesian multilevel models (Bürkner, 2017). However, even without this feature, this pipeline gives any user the ability to analyse their own data, and the flexibility to incorporate additional approaches and analyses.

## Acknowledgements

DP was funded by a NERC standard grant and by the University of Exeter and SP was funded by NERC grants (NE/S000348/1 and NE/M020843/1).

## Author contributions

DP conceived the ideas and designed the pipeline. DP authored the *R* package and wrote the initial draft. All authors contributed to developing the manuscript and gave final approval for publication.

## Supplementary Information

**Table 1:**
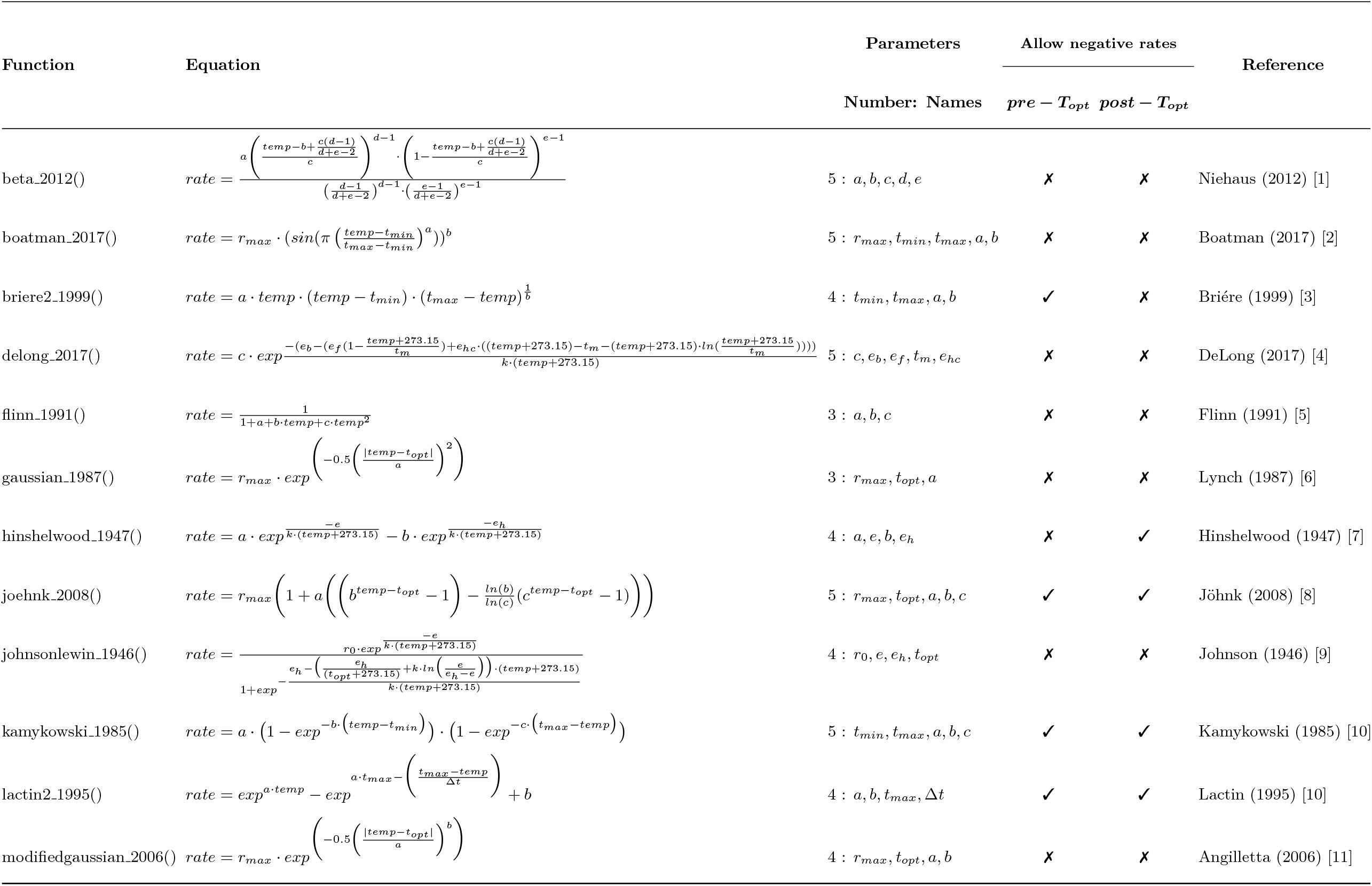
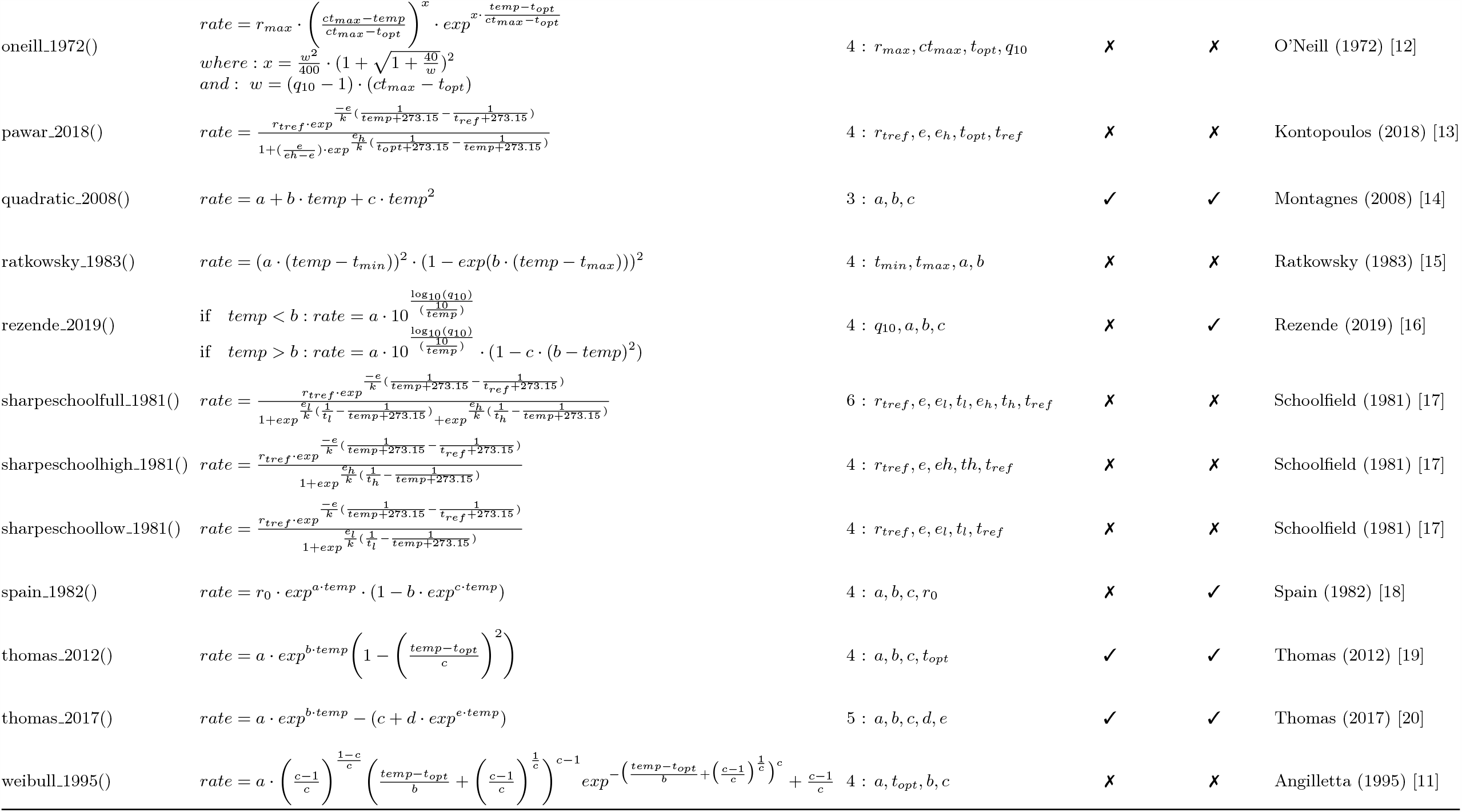
Summary of equations available in rTPC

## Notes

https://padpadpadpad.github.io/rTPC/

### Competing Interest Statement

The authors have declared no competing interest.

